# Comparison of Biofilm quantification in strains of *Candida auris* and *Candida albicans* evaluated by means of Crystal violet, MTT, ATP and NBTZ/BCIP Assays

**DOI:** 10.64898/2026.03.15.711842

**Authors:** Jens Jacob

## Abstract

The study presented here shows Biofilm quantification in microtiter plates in strains of *Candida auris* and *Candida albicans* evaluated by means of Crystal violet, MTT, ATP-Luminescence and NBTZ/BCIP assays. The results showed significant differences in biofilm formation between *Candida auris* and *Candida albicans* but also within *Candida auris* outbreak strains in contrast to *Candida auris DSM 21092* reference strain.

## Background

Yeasts of the genus *Candida* are members of the *Metschnikowiaceae* and they are phylogenetically closely related to the human non-pathogenic *Saccharomycetacea*. Despite the fact that DNA homology is high between *Saccharomycetacea* and *Metschnikowiaceae*, i.e. only the genera *Candida* within the *Metschnikowiaceae* shows human pathogenic capabilities. Besides the previously known *Candida albicans* there is currently a rise in papers mentioning *Candida auris* as a new emerging human pathogenic yeast [6,16]. *Candida* (and hereby also *C. auris*) are now known worldwide as a cause of human infections, especially in immunocompromised patients. Due to its increasing resistance against antifungal drug [21,23]. However, *Candida* are also able to survive several days under certain environmental conditions especially and foremost in highly regulated biofilms [8]. They also quickly develop resistance against surface disinfectants [7,9,10].

However, in contrast there is both an increasing number of sample submissions to the German consular laboratory for mycoses NRZ Nationales Referenzzentrum für Invasive Pilzinfektionen (https://www.nrz-myk.de/home.html) [13] and to the German Infectious Diseases Reporting System for infectious disease resulting in a continuous rise of reported *Candida auris* cases (https://meldung.demis.rki.de/portal/shell/#/welcome; https://survstat.rki.de/Default.aspx) [14].

That has been probably caused by the higher alert preparedness of practical doctors since establishing the law for the duty of reporting of putative cases of invasive candidiasis in 2022. However, the altered efficacy of disinfectants used in current hospital hygiene practice could also be a reason for changes in observation of biofilms formation capacity. The synthesis and excretion of such altered biofilm matrices and their role in pathogenicity towards humans had been previously described and the applied methods were compared [1, 2, 4, 20, 22].

Various methods to quantify biofilm formation capacity for *Candida albicans* were used in the current study according previously described [8]. Biofilms grown in 96 well plates and quantified by means of Crystal violet, MTT, ATP and Alkaline Phosphatase assays we used to evaluate both biofilm formation between *C*.*auris* and *C*.*albicans* and to suitability of the different assays used in this study.

## Material and Method

### Yeast strains

We investigated in this study a) standard reference strains of *C. auris* DSM 21092 [17], *C. auris* DSM 105986, b) nosocomial outbreak strains *C. auris Charité* 588, 607, 610, 611 as described in paper [11] and c) *C. albicans* MH 203 wild type and genetically modified gene deletion mutants *C. albicans* 527 *ΔΔals1als3* and *C. albicans* 485 *Δals3* [19]. The genetically modified *Candida albicans* strains were genetically defective for the *als1als3* adhesine genes. *Candidae* defective in adhesine genes are usually known to show reduced biofilm development potential [23]. All *Candida* strains generally grown and handled according DIN/EN 17387 in analogy as to be described there for *C. albicans* in the respective norm. For the strains described above throughout the text and figure’s respectively the following abrevations were use: *C. auris Charité* 588, 607, 610, 611 (CHAU CHAR 588, 607,610,611), *C. auris* DSM 21092 [17], *C. auris* DSM 105986 (CAU DSM 21092, CAU DSM 105986), *C. albicans* MH 203, *C. albicans* 527 *ΔΔals1als3, C. albicans* 485 *Δals3* (CALB 203wt, CALB 527 ΔΔ, CALB 485 Δ).

### Biofilm cultivation

Prior to any biofilm quantification measurement biofilms were cultivated in 96-well microtiter plates over night at 30° in a TECAN 200 M Pro multi-well shaker under medium orbital shaking with the following plate setup:

For all assays performed here each strain in each dilution row (−1 until -4 dilution, corresponding to 10^7^ until 10^4^ CFU/ml) was inoculated in three wells for technical triplicates and additionally, each 96 well plate contained 50 µl of the respective diluted sample. Selected wells were also filled with water as negative controls for subsequent quantification experiments. Then, as an follow up to this prior biofilm measurement biofilm formation growth period (to allow regular biofilm formation before any test), microtiter plates were further processed to quantify biofilm formation measured by means of the 5 different assays used in this study and described here in material and methods section in detail.

The CFU/ml values in the figures were obtained from plating one out of three technical replicates after biofilm growth prior quantification measurements.

### Biofilm quantification methods

#### Crystal violet assay

For biofilm quantification by means of Crystal Violet the Assay kit from Abcam (art nr. 232855) was used. After biofilm growth the supernatant containing planktonic cells was removed, and then to each well 50 µl Crystal violet solution was added for 20 min. After this incubation, the crystal violet staining solution was removed and the wells were 5 times washed in water. After overnight drying 200 µl Methanol was added to each well. Again after 20 min incubation the so released crystal violet washed out from the biofilm was measured both at absorbance OD 570 nm. Because one sample measured consisted of three technical replicates each one of these was not stained but instead got plated on MEA Agar for CFU/ml estimation to be used for comparison in the respective graphics. That step was done identical also in the other biofilm quantification assays. The exact number of biological/technical replicates investigated is given for each assay in Sup. Files 1. Tab.2.

#### MTT (OD 570 nm) Assay

The 3-(4,5-dimethylthiazol-2-yl)-2,5-diphenyltetrazolium bromide (MTT) based assay Cye Quant MTT Cell Proliferation Assay (art nr. v13154) from Thermofisher (Invitrogen) is a widely used colorimetric method for assessing cell viability, proliferation, and cytotoxicity. Briefly 96 well multi-well dishes were inoculated with 10^-1^-10^-4^ dilutions of overnight cultures obtained from 8 *Candida* strains each in triplicates. The strains in the multi well-dishes were grown for overnight each at 30 °C under orbital shaking in a 200 M Pro Multi-well reader (TECAN). For each experiment two 96 well multi well dishes were inoculated and used. The biofilm containing dish was so further proceeded for the MTT measurement assay by adding 10 ul MTT to each well. Afterwards 100 ul lysis solution (SDS in 1N HCL) was added and after 4 hrs lysis incubation again the MTT formation was measured at OD 570 nm at the TECAN multi well reader.

For more realistic quantitative interpretation of the MTT OD570 (discrimination of LIVE/DEAD biofilms) results the assay was refined by addition of an inhibiting antibiotic substance, here in case of *Candida* 300,100,30,10, 0 µg/ml Nourseothricin (corresponding to 30,10,3,1,0 µl Nourseothricine solution in Fig.5, Tab.1) to the respective dilution row of the respective triplicates within this new **OD 570 Nourseothricin inhibition Assay**. The so obtained OD570 (Nourseothricin treated) values reflects the respective signal presence in growth inhibited *Candida* cells. Subtracting these OD570 (Nourseothricin treated) from the not-Nourseothricin treated “LIVE” OD 570 values, gives an (*Candida* strain specific) quota indicating the relation of metabolic alive cells showing MTT signal in contrast to lesser pronounced signal presence in cells growth inhibited by Nourseothricin. (Quota (strain specific) = OD 570 (LIVE)-OD570 (Nourseothricin treated, Tab. 1).

**Table 1.** OD 570 MTT. Nourseothricin inhibition Assay OD 570 MTT Nourseothricin inhibition Assay OD 570 MTT inhibition Assay. The so obtained OD570 (Nour) values reflects the respective formazan formation in growth inhibited *Candida* cells only. Subtracting these OD570 (Nourseothricine treated) from the non-Nourseothricine treated OD 570 values gives an (candida strain specific) quota indicating the relation of MTT signal formation in contrast to MTT signal formation in the Nourseothricin growth inhibited *Candida cells*. Quota (strain specific) = OD 570-OD570 (Nour).

| Candida Strain | Dilution | CFU/ml | ΔOD570-OD 570 (NOUR) Quota | OD 570 (NOUR) | Number of experiments (as technical triplicates each) |
| --- | --- | --- | --- | --- | --- |
| **CAU CH 588** | **1/10** | $10^7$ | **1.2** | **1.6** | **5** |
| **CAU CH 611** | **1/10** | $10^7$ | **1.3** | **1.8** | **5** |
| CALB $\Delta\Delta$527 | 1/10 | $10^7$ | 0.6 | 2.2 | 5 |
| **CAU DSM 21092** | **1/10** | $10^7$ | **1.4** | **1.8** | **5** |
| **CAU CH 607** | **1/10** | $10^7$ | **1.6** | **1.8** | **5** |
| CALB Δ485 | 1/10 | $10^7$ | 0.7 | 1.6 | 5 |
| CALB 203wt | 1/10 | $10^7$ | 0.7 | 1.8 | 5 |
| **CAU CH 588** | **1/100** | $10^6$ | **0.5** | **0.4** | **5** |
| **CAU CH 611** | **1/100** | **10<sup>6</sup>** | **0.6** | **0.4** | **5** |
| CALB $\Delta\Delta 527$ | 1/100 | 10<sup>6</sup> | 0.1 | 0.6 | 5 |
| **CAU DSM 21092** | **1/100** | **10<sup>6</sup>** | **1.3** | **0.5** | **5** |
| **CAU CH 607** | **1/100** | **10<sup>6</sup>** | **1.0** | **0.4** | **5** |
| CALB $\Delta 485$ | 1/100 | 10<sup>6</sup> | 0.4 | 0.4 | 5 |
| CALB 203wt | 1/100 | 10<sup>6</sup> | 0.4 | 0.5 | 5 |
| CAU CH 588 | 1/1000 | 10<sup>5</sup> | 0.2 | 0.2 | 5 |
| CAU CH 611 | 1/1000 | 10<sup>5</sup> | 0.15 | 0.14 | 5 |
| CALB $\Delta\Delta 527$ | 1/1000 | 10<sup>5</sup> | 0.12 | 0.19 | 5 |
| CAU DSM 21092 | 1/1000 | 10<sup>5</sup> | 0.2 | 0.2 | 5 |
| CAU CH 607 | 1/1000 | 10<sup>5</sup> | 0.04 | 0.17 | 5 |
| CALB $\Delta 485$ | 1/1000 | 10<sup>5</sup> | 0.02 | 0.15 | 5 |
| CALB 203wt | 1/1000 | 10<sup>5</sup> | 0.04 | 0.18 | 5 |

As precondition for that new Nourseothricin inhibition assay all *Candida* strains used in this study were tested susceptible against 30µg/ml Nourseothricine (Fig. 5). The terminus Nourseothricine is used throughout now in this text despite of the fact that “Nourseothricine” consists of, in analogy to the terminus “Polymyxine”, always of a mixture of different streptothricines with different side chain length’s (as results of manufacturing process) [19].

### ATP Assay/Determination of intracellular ATP content

ATP-Luminescence was measured with BacTiter-Glo® Microbial Viability Assay (Promega, cat. nr. G8231, Madison, WI, USA). All experiments were done in white-, 96 well-microplate dishes designed for luminescence measurements (Thermoscientific, cat. nr. 236105, Waltham, MA, USA).

The BacTiter-Glo ^®^ reagent was used according manufacturer’s manual.

For the setup of the experiments, the respective *Candida* strains were diluted 10fold from -1 until -4 in a standardized buffer system and seeded in triplicates in the wells of the respective 96 well dish and incubated over night at medium orbital shaking at 200 rpm at 30°C for the pre-measurement biofilm growth period.

Then, after adding of 25ul BacTiter-Glo^®^ working solution the plate covered with Adhesive Film ® (NEOLAB, cat. nr.7-5170). Aerobic incubation was performed until 18hrs and ATP formation continuously monitored up to 18hrs at time points 0min, 15min and 18hrs. 15 min values were used for results interpretation and presented in this study. In the time periods indicated here, the bacteria were lysed and ATP was finally measured via the luminescence-reaction via the TECAN Infinite 200 M Pro (Tecan, Gröding, Austria) reader and expressed as relative light units (RLU) as described in the manual of the BacTiter-Glo^®^Kit (BacTiter-Glo^®^ Microbial Viability Assay (Promega, cat. nr. G8231, Madison,WI, USA). That was as outlined in detail previously [12]. For that the “standard-automatic” luminescence measurement program was used.

Colony forming units (CFU) were determined in parallel by plating one triplicate out of three on MEA agar each to which no lysis mix was added. They assay surviving CFU’s were counted after 2 days on. All luminescence experiments were done at least as three biological separate experiments.

### NBTZ/BCIP Assay (Indigo formation)

The detection of endogenous alkaline phosphatase the AP Conjugate substrate Kit (Bio-Rad, cat. Nr. 1706432, Hercules, CA, USA) was performed according manufacturer’s manual. All experiments were done in black, transparent-, 96 well-microplate dishes designed for fluorescence measurements (Thermoscientific cat. nr. 165305, Waltham, MA, USA). For the setup of the experiments, the respective Candida strains were diluted 10fold from -1 until -4 in standardized buffer system and seeded in triplicates in the wells of the respective 96 well dish and incubated over night at orbital shaking at 200 rpm at 30°C for the pre-measurement biofilm growth as already outlined here.

After adding of 50µl NBTZ/BCIP (Nitroblue Tetrazolium/5-Bromo-4-chloro-3-indolyl Phosphate) working solution aerobic incubation was performed again until 18hrs and indigo formation was continuously monitored all these 18hrs. Values obtained at 0min, 15min and 18hrs were used for results interpretation. In this time period, the cells were lysed and indigo formation was measured via fluorescence detection at an TECAN Infinite 200 M Pro (Tecan, Gröding, Austria). For indigo formation measurements were done at OD 620nm.

Colony forming units (CFU) were determined in parallel by plating the 3^rd^ triplicate on MEA agar. They were counted after 2 days on. All experiments were done at least three times.

### Statistical analysis

Graph Pad Prism 7.04 (San Diego, USA) was used for statistical analysis (Mann-Whitney-U test) of the data.

Data of the biofilm growth assays were based on at least three independent experiments using triplicates each. For crystal violet measurements 6 biological replications were run, 4 for ATP measurement and both 3 for ATP and NBTC/BCIP (indigo formation) assays. The respective p-values, geometric means, standard derivations and the belonging 95% confidence interval values were not shown in the results figures but given in Supplemental information File 1 (Tab. 2)

## Results

The results obtained for the crystal violet measurements at 570nm OD mode showed (Fig.1) a statistically significant, different yield of crystal violet in outbreak strain CAU CHAR 607 in contrast to CAU DSM 21092 and CAU DSM 105986 reference strains, as well to *C. albicans 527ΔΔals1als3* (adhesine defective) double mutant [19] and finally with respect the outbreak strains CAU CHAR 611 and CAU CHAR 588.

**Fig. 1.**
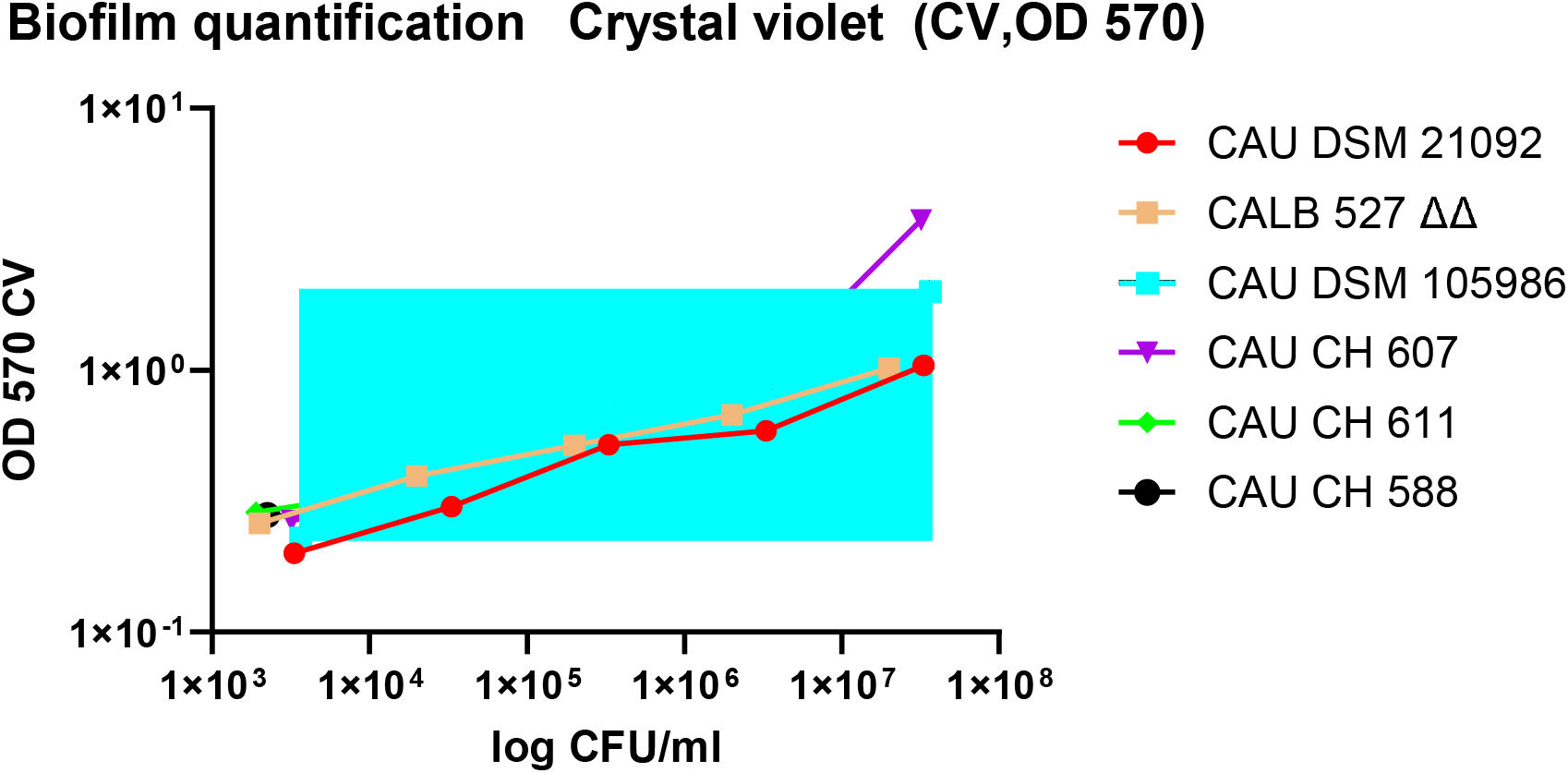
CV OD 570. *In*-*vitro* survival of various *Candida* species in a 96 well based, overnight biofilm development set-up, determined by Crystal violet measurements (CV, OD 570nm) and CFU plating after various times of exposition. P-Values (Significance Y/N) CAU DSM21092/CAU CH 607 10 fold diluted p<0.0001 Y CV OD 570; CAU DSM21092/CAU CH 607 100 fold diluted p<0.0001Y CV OD 570; CAU DSM21092/CAU CH 607 1000 fold diluted p 0.0176 Y CV OD 570; CAU CH 607/CAU CH 588 10 fold diluted p<0.0001 Y CV OD 570; CAU CH 607/CAU CH 611 10 fold diluted p<0.0001 Y CV OD 570; CAU DSM21092/CAU DSM 105986 10 fold diluted p 0.01 Y CV OD 570; CAU DSM21092/CAU CH 588 10 fold diluted p<0.0001 Y CV OD 570 The respective standard deviations and confidence interval values are listed in Sup. Information File 1.

The results obtained for the MTT measurements at 570nm OD mode (Fig.2) were monitored after 4 hrs (recommended by manufacturer’s instructions for use of the kit). The respective data obtained gave statistically significant differences (Fig. 2). This is valid for the 100, 1000, 10000fold dilutions of the initial cell stock (10^5^, 10^4^ resp. 10^3^ cells per well) a grouping of strains with statistically significant MTT values, i.e. a) CAU CHAR 607/ CAU DSM 21092 and b) CAU CHAR 588/611 was observed (Sup. File 1, Tab. 2). Within the clinical isolates CAU CHAR 607/611/588 the strain CAU CHAR 607 showed a significant higher (comparable to CAU DSM 21092) MTT content in contrast to strains CAU CHAR 611/588.

**Fig. 2.**
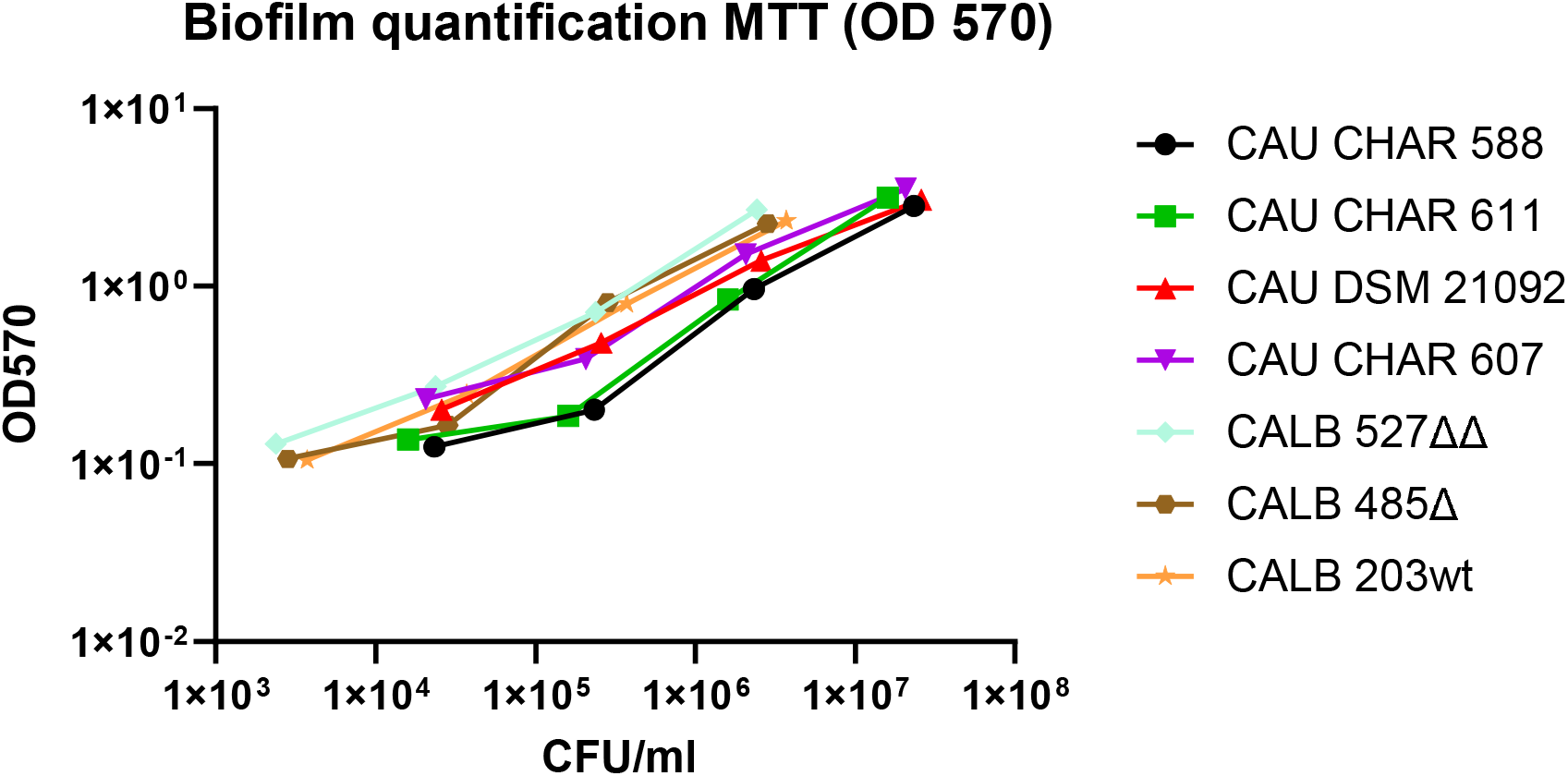
MTT OD 570. *In*-*vitro* survival of various *Candida* species in a 96 well based, overnight biofilm development set-up, determined by Dimethyl(thiazol-2-yl)-2,5-diphenyltetrazolium bromide (MTT) measurements (OD 570) and CFU plating after various times of exposition. P-Values (Significance Y/N) CAU DSM21092/CAU CH 588 100 fold diluted p 0.01 Y MTT (OD 570); CAU DSM21092/CAU CH 611 100 fold diluted p 0.0006 Y MTT (OD 570); CAU CH 607/CAU CH 611 100 fold diluted p<0.0001 Y MTT (OD 570); CAU CH 607/CAU CH 588 100 fold diluted p 0.0003 Y MTT (OD 570); CAU CH 607/CAU CH 588 1000 fold diluted p 0.028 Y MTT (OD 570); CAU CH 607/CALB CH Δ485 1000 fold diluted p 0.001 Y MTT (OD 570); CAU DSM 21092/CALB Δ485 1000 fold diluted p<0.0001 Y MTT (OD 570); CALB 203 wt /CALB Δ485 1000 fold diluted p 0.006 Y MTT (OD 570) The respective standard deviations and confidence interval values are listed in Sup. Information File 1.

Summing up, identical strains now measured in MTT test (Fig.2) in contrast to results of crystal violet testing (Fig.1) did not show the differences between CAU CHAR CH 607 and CAU DSM 21092 as shown in Fig.1. The ratio behind that finding may be that crystal violet staining is mostly directed towards Polysaccharides, Glucanes whereas MTT staining corresponds to Formazan accumulation in the respiratory chain of the mitochondria, i.e. MTT formation could be considered more as a LIVE/DEAD indicator [3] despite of the point similar that similar cell numbers in biofilms were present in both assay setups.

Hence, to get more insight into the discrepancy between Fig1 and Fig. 2, i.e. the question will be metabolic active resp. inactive parts in crystal violet measurements the Nourseothricin inhibition assay was performed.

The results of the MTT Nourseothricin inhibition assay (Tab. 1, Fig. 5) showed significant different quota between both *Candida auris* and *Candida albicans* strains only with respect to the dilution of the initial stock. However, and most importantly to note with respect to the general applicability of such assays: Even under growth inhibiting conditions significant great amounts of MTT can be measured, i.e. even dead cells show significant amounts of MTT formation. This has to be taken in consideration with respect to the general interpretation of such biofilm quantification assays, i.e. only a part of the measured, cell metabolism based, new formation of MTT may be connected to the metabolic activity of the just set up assay, a bigger part is probably linked to the polysaccharide made biofilm matrix.

So here we demonstrated, that the so adjusted assay shows, even then the cells are dead, significant measured values for the presence of the MTT based signals. Because of that, the application of the described formula: Quota (strain specific) = OD 570-OD570 (Nourseothricine inhibited) seems to be a valuable improvement for the estimation of the real biofilm matrix formation with respect to the cell dilution used in experiment.

Hence necessary to explain, that the inhibition of *Candida auris* and *Candida albicans* by Nourseothricin was not only shown by the MTT OD 570 results but also by plating the respective, Nourseothricin treated cells on MEA agar (Fig.5). The *Candida auris* strains showed an average log 6 reduction, the *Candida albicans* strains a respective log 3 reduction at 30 µg/ml in the MHK test applied (Fig. 5).

The so surprisingly obtained finding of the inhibitory effect of the “old known” 2^nd^ generation aminoglycoside antibiotic Nourseothricine (mixture of streptothricines A-F with different side chain length) will be addressed in a further study, because the effect is worth to investigate because of the general rise of antibiotic/antifungal resistance of *Candida* species.

### ATP Assay

The ATP content of the strains used in this study was monitored for the initial seed (0 minutes) 15 minutes (recommended by manufacturer instructions for use of the kit) and at 18 hrs respectively. Because the results at 18 hrs showed only an overall, statistically not significant, ATP decline for all strains the data are not shown in detail here. In contrast the data obtained at 15 min (use recommended also according manufacturer instructions) gave statistically significant differences (Fig. 3). As for the 100 resp. 1000fold dilutions of the initial cell stock (10^5^ resp. 10^4^ cells per well) a grouping of strains with statistically significant ATP values, i.e. a) CAU CHAR CH607/ CAU DSM 21092, b) CAU CHAR CH 588/611 and c) CALB 527ΔΔ, 485Δ, 203wt, was observed (Sup. File 1, Tab. 2). Of the clinical isolates CAU CHAR 607/611/588 the strain CAU CHAR CH 607 showed a significant higher (comparable to CAU DSM 21092) ATP content in contrast to strains CAU CHAR 611/588. Both these groups showed a contrast to group c), the reference strains C. albicans 527ΔΔ, 485Δ, 203wt used in this study. So from that viewpoint of cell ATP content we conclude that the strains investigated here could be grouped in these respective three groups (a, b, c).

**Fig. 3.**
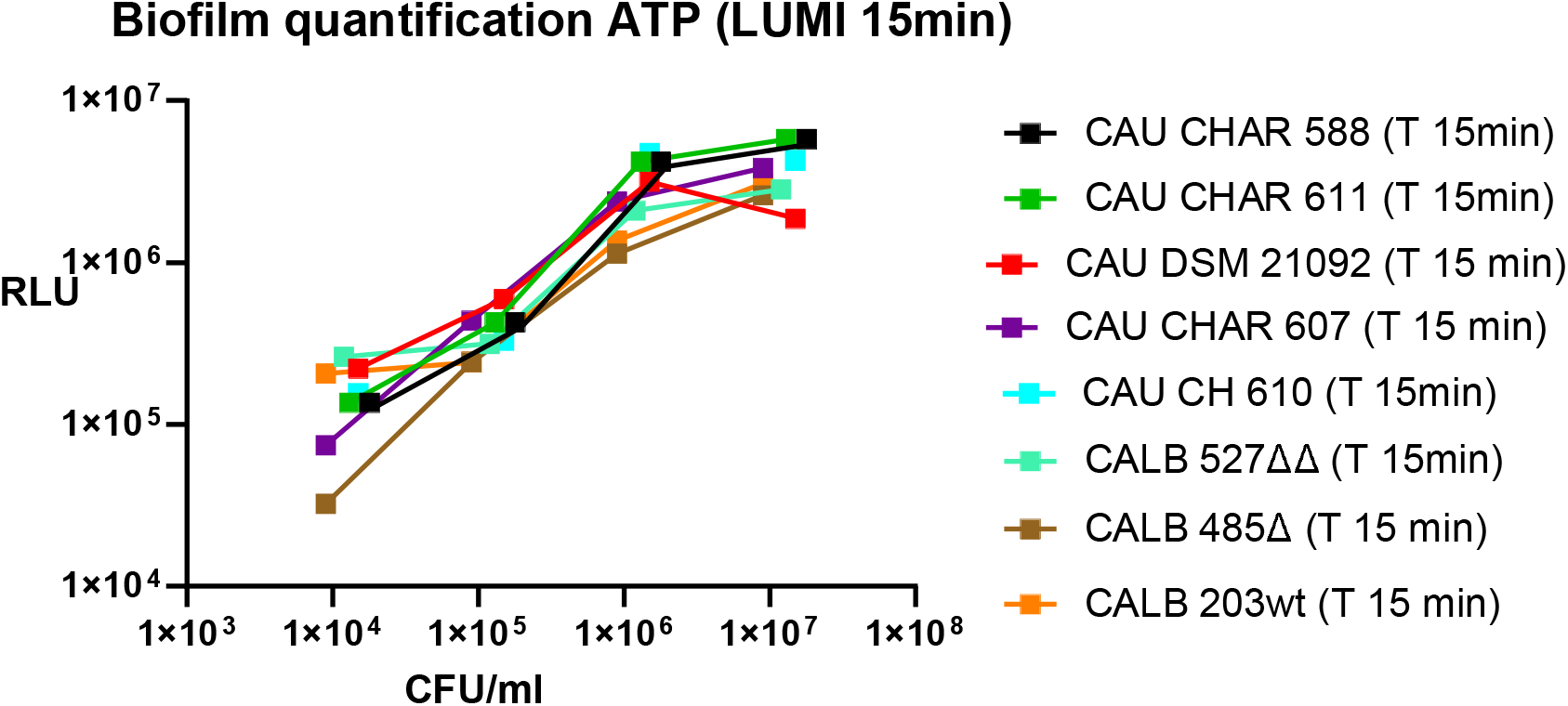
ATP Assay. *In-vitro* survival of various *Candida* species in a 96 well based, overnight biofilm development set-up, determined by ATP Luminescence Assay and CFU plating after various times of exposition. P-Values (Significance Y/N) CAU DSM21092/CAU CH 611 LUMI -10 fold diluted 15 min p 0.0003 Y; CAU CH 607/CAU CH 588 LUMI-10 fold diluted 15min p 0.05 Y; CAU CH 607/CAU CH 588 LUMI-100 fold diluted 15min p<0.0001 Y; CAU CH 607/CALB Δ485 LUMI-10 fold diluted 15 min p 0.02 Y; CAU CH 607/ CALB Δ485 LUMI-100 fold diluted 15 min p 0.003 Y; CAU CH 588/CALB Δ485 LUMI-10 fold diluted 15 min p<0.0001 Y; CAU CH 588/ CALB Δ485 LUMI-100 fold diluted 15 min p<0.0001 Y; CAU DSM21092/CALB ΔΔ527 LUMI - 100 fold diluted 15 min p 0.03 Y; CAU DSM21092/CALB Δ485 LUMI -100 fold diluted 15 min p 0.02 Y; CAU DSM21092/CALB 203wt LUMI -100 fold diluted 15 min p 0.004 Y; CAU CH 607/CALB ΔΔ527 15 min p<0.0001 Y; CAU CH 607/CALB 203wt LUMI-100 fold diluted 15 min p 0.007 Y; CAU DSM21092/CALB Δ485 LUMI -1000 fold diluted 15 min p 0.002 Y; CAU CH 607/CAU CH Δ485 LUMI-1000 fold diluted 15 min p 0.03 Y; CAU CH 607/CALB 203wt LUMI-1000 fold diluted 15 min p 0.01 Y; CALB ΔΔ527/CALB 203wt LUMI-100 fold diluted 15 min p 0.004 Y; CALB ΔΔ527/CALB Δ485 LUMI-100 fold diluted 15 min p 0.0001 Y The respective standard deviations and confidence interval values are listed in Sup. Information File 1.

#### NBTZ/BCIP Assay

(Nitroblue-Tetrazoliumchloride) /BCIP (5-Brom-4-chlor-3’-indolylphosphat p-toluidine salt)

The combination of NBT (Nitroblue-Tetrazoliumchloride) and BCIP (5-Brom-4-chlor-3’-indolylphosphat p-toluidine salt) gives an intense, un-soluble black violet precipitation (Indigo blue), if it reacts with alkaline (here the endogenous in the yeast) Phosphatase. So here the Indigo formation (measured via endogenous alkaline phosphatase activity) of the strains was then monitored for the initial seed (0 minutes), 15 minutes and at 18 hrs respectively (to measure at the same time points as it was done in the ATP assay). Because the results at 18 hrs showed only an overall, statistically not significant, Indigo signals rise for all strains, the resulting data for 18 hrs are not shown here. In contrast the data obtained at 15 min showed statistically significant differences (Fig. 4). So valid for the 10, 100 resp. 1000fold dilutions of the initial cell stock (10^6^, 10^5^ resp. 10^4^ cells per well) a grouping of strains with statistically significant ATP values, i.e. a) CAU CHAR 607/ CAU DSM 21092, b) CAU CHAR CH 588/611 and c) CALB 527ΔΔ, 485Δ, 203wt, was observed (Sup. File 1, Tab. 2). Of the clinical isolates CAU CHAR 607/611/588 the strain CAU CHAR 607 showed a significant higher (comparable to CAU DSM 21092) Indigo (Formazan) content in contrast to strains CAU CHAR 611/588. Both these groups showed a contrast to group c, the reference strains CALB 527ΔΔ, 485Δ, 203wt used in this study. From viewpoint of, indigo measurement based, endogenous cell alkaline phosphatase activity, we conclude also that the strains investigated could be grouped in these three groups.

**Fig. 4.**
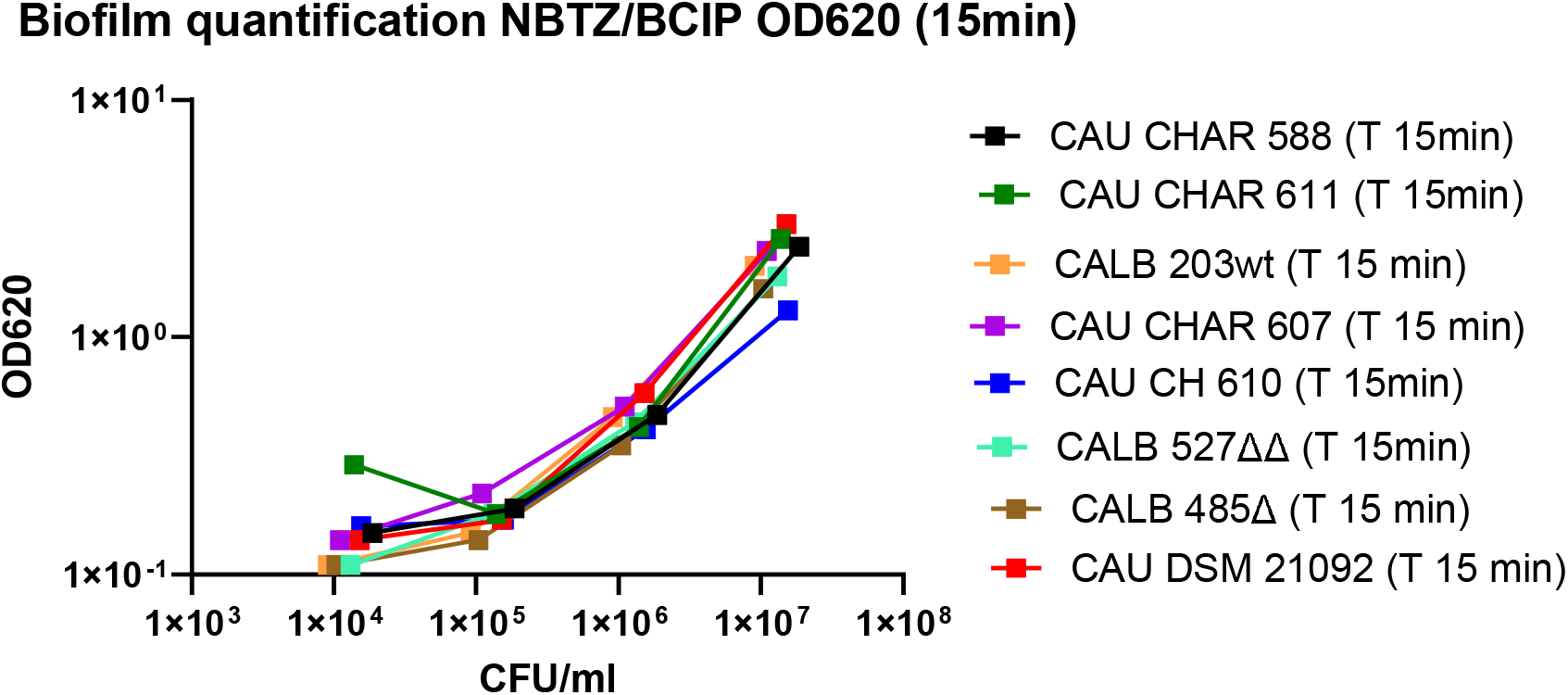
NBTZ/BCIP Assay (Indigo formation) *In*-*vitro* survival of various *Candida* species in a 96 well based, overnight biofilm development set-up, determined by an NBTZ/BCIP (Formazan-Indigo formation) endogenous alkaline phosphatase assay and CFU plating after various times of exposition. P-Values (Significance Y/N) CAU CH 588/CALB CH Δ485 NBTZ/BCIP-10 fold diluted 15 min p<0.0001 Y; CAU CH 588/CALB 203wt NBTZ/BCIP-10 fold diluted 15 min p<0.0001 Y; CAU CH 21092/CALB Δ485 NBTZ/BCIP-10 fold diluted 15 min p<0.0001 Y; CAU CH 607/CALB ΔΔ527 NBTZ/BCIP-10 fold diluted 15 min p<0.03 Y; CAU CH 607/CALB Δ485 NBTZ/BCIP-10 fold diluted 15 min p 0.004 Y; CAU DSM21092/CALB Δ485 NBTZ/BCIP-100 fold diluted 15 min p 0.0007 Y; CAU CH 607/CALBΔ485 NBTZ/BCIP-100 fold diluted 15 min p 0.003 Y; CAU CH 21092/CALB Δ485 NBTZ/BCIP -1000 fold diluted 15 min p<0.0001 Y; CAU CH 607/CALB CH Δ485 NBTZ/BCIP-1000 fold diluted 15 min p 0.0001 Y; CAU CH 588/CALB Δ485 NBTZ/BCIP-1000 fold diluted 15 min p 0.001 Y; CAU CH 611/CALB Δ485 NBTZ/BCIP-1000 fold diluted 15 min p 0.01 Y; CAU CH 21092/CAU CH 607 NBTZ/BCIP-1000 fold diluted 15 min p 0.007 Y; CALB 203wt/CALB Δ485 NBTZ/BCIP-100 fold diluted 15 min p 0.03 Y; CALB 203wt/CALB Δ485 NBTZ/BCIP-10 fold diluted 15 min p<0.0001 Y; CALB 203wt/CALB ΔΔCH 527 NBTZ/BCIP-10 fold diluted 15 min p<0.004 Y The respective standard deviations and confidence interval values are listed in Sup. Information File 1 (Tab. 2).

The quite comparable results of both the ATP Assay and NBTZ/BCIP Assay (Indigo/Formazan formation) could be explained because the formation of Indigo depends on the ATP content of the cell, i.e. if Indigo gets build up the cell depletes the cells ATP reservoir. Hence both assays measured the same cellular process, however, according our knowledge, that was not shown in that way in previous studies in this field of medical microbiology. Worth to note that the ATP demanding formation of Indigo blue via NBTC/BCIP colour reaction is used as indicator for cellular endogenous alkaline phosphatase activity.

The (maximal) 5 bars for each organism represents application of 30µl, 10μl, 3µl, 1μl, 0µl Nourseothricine each. All this information would not fit as text in the legend of the y-axis. In Fig. 5 for each strain only STR 30µl is written indicating the first bar of maximum five. If a bar is missing it means no *Candida* growth was measured by plating.

**Fig. 5.**
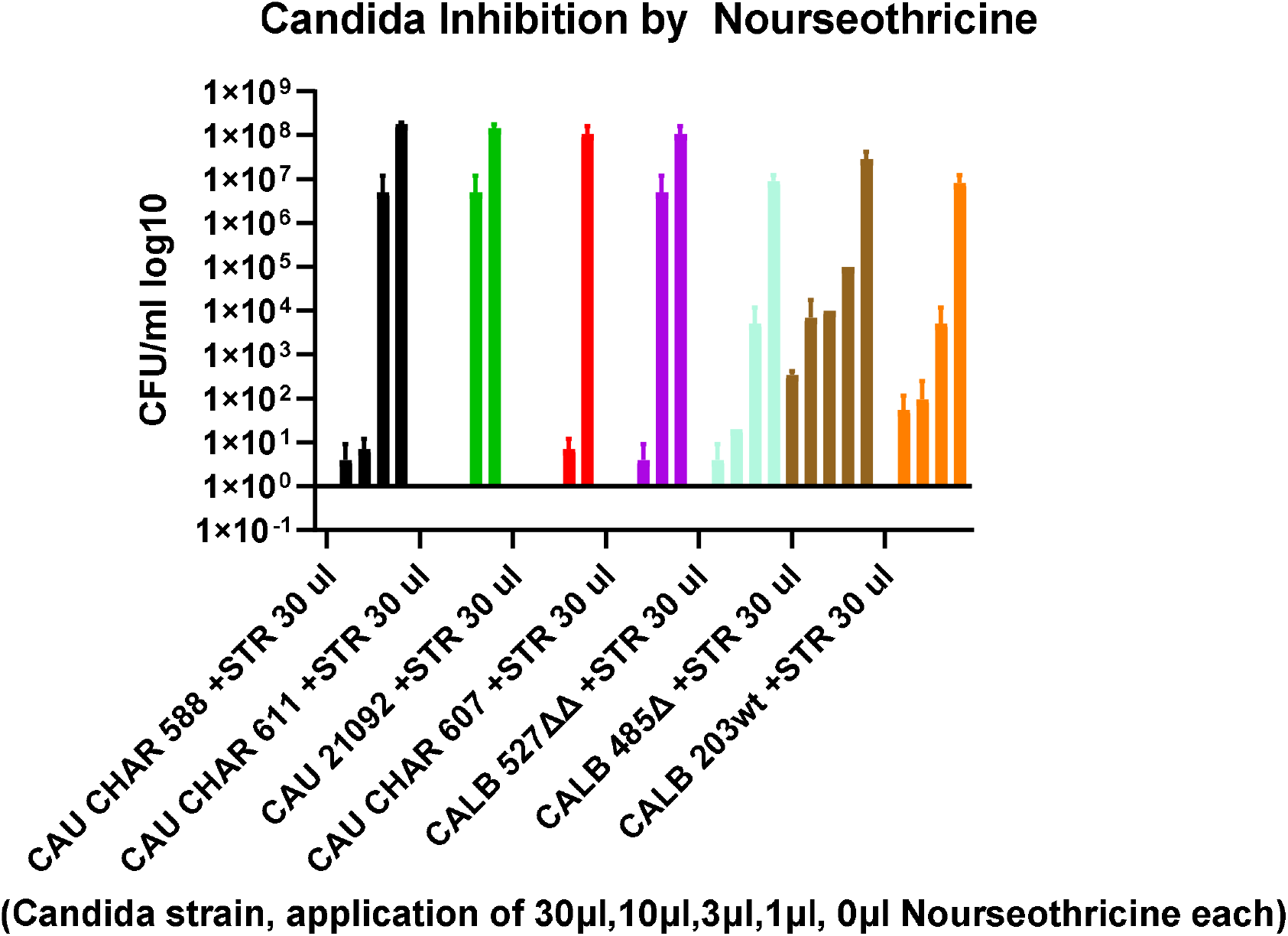
Nourseothricine Inhibition Assay of *Candida* strains. Inhibition of *Candida* by Nourseothricine The figure shows the *Candida* strains, for each strain the CFU’s (in bars from left) after application of 30µl, 10µl, 3µl,1µl, 0µl Nourseothricine each. Only the bars (corresponding to the Nourseothricine concentration used) which showed CFU’s after plating, were displayed.

If we have here for example two red bars for strain CAU DSM 21092 only, it means the 30,10 and 3µl Nouseothricine solution inhibited the strains totally (no CFU’s detectable) whereas for 1µl and 0 µl (growth control) growth was visible. According that scheme the graph could be read for all strains.

The description of y axis (30,10,3,1,0 µl Nourseothricine solution in Fig.5, Tab.1) corresponds to300,100,30,10, 0 µg/ml Nourseothricin each.

## Discussion

The results presented in this study, obtained by 5 different methodological approaches, i.e. Crystal violet OD 570, OD570 MTT/ Nourseothricin inhibition, ATP measurement and NBTZ/BCIP ((Indigo formation, caused by endogenous alkaline phosphatase) detection assay, showed a different degree of biofilm formation not only between *Candida auris* and *Candida albicans* but also between *Candida auris* strains from an outbreak [11] in comparison to *Candida auris* reference strain *DSM 21092. Candida auris* outbreak strain *607* showed quite comparable biofilm formation characteristics as *Candida auris* reference strain *DSM 21092*. However, only the Crystal violet assay OD570 showed significant differences between *Candida auris* outbreak strain CHAR *607* and *Candida auris* reference strain *DSM 21092*.

The results for the **MTT, ATP** and **NBTZ/BCIP Assay (Indigo formation)** respectively, showed a grouping of strains with slightly other statistical significance, i.e. a) CAU CHAR 607/ CAU DSM 21092, b) CAU CH 588/611 and c) CALB 527ΔΔ, 485Δ, 203wt.

Interestingly the grouping in the newly introduced, **Nourseothricine inhibition assay** (Tab.1.) showed the same grouping with respect to the described quotas i.e. a) CAU CHAR 607/ CAU DSM 21092, b) CAU CH 588/611 and c) CALB 527ΔΔ, 485Δ, 203wt.

**In summary**, the selection of biofilm quantification methods used in this study was found not only useful for biofilm quantification as itself, but also applicable for further research with respect to epidemiological information about circulating strains. In addition to the already previously introduced CV, MTT, ATP and NBTZ/BCIP assay’s the new **Nourseothricine inhibition assay** is a new promising tool because here first time the results were obtained from a relation derived not only from signals from alive cells but also from dead cells.

### Epidemiology

For evaluation of the biofilm quantification results in context to the epidemiological knowledge what we have about the strains we compared in this study, it is to remark that CAU CHAR 588 is the immediate descendant of the index strain in the *C*.*auris* outbreak description published [11]. Strain CAU CHAR 607 is a follow-up isolate of a contact person within the same outbreak. However, CAU CHAR 611/610 were isolated from an, short time later recognized, follow up outbreak in the same medical facility.

The identity of these both strains can be seen in Fig.3 with respect to the ATP and in Fig. 4 with respect to the Indigo formation.

For the reference strains: CALB 203 wt is an outbreak type strain and CALB 485Δ, CALB 527ΔΔ are descendants created by genetical manipulations i.e.by locus specific deletion mutagenesis of the Δ*als1* respectively ΔΔ*als1als2* adhesivity genes. Both they play a prominent role for in vitro and in vivo biofilm formation capacity [19]. Our observed grouping of the strains a) CAU CH607, CAU DSM 21092, b) CAU CHAR 588/611 and c) CALB 203wt, 485Δ, 527ΔΔ goes so far round with that. However, according our findings, CAU CH 607 shows significant different biofilm formation potential as international reference strain CAU DSM 21092 shows it. Also, because strains CAU CHAR 588, 611 have identical capabilities we assume that is that one outbreak, i.e. the described 2^nd^ outbreak [11] was more a persistent “post outbreak” circulation of the strains from outbreak one. The *Candida albicans* strains are clearly different. These different capabilities according to the respective mutations, were also shown to be significant (Sup. File 1, Tab.2). All this leads to our conclusion that the sensitivity of the previous 4, now 5 methods used here were quite sufficient for the purpose of biofilm quantification as done in this study.

## Supporting information

Sup File 1 Standard deviations and confidence intervals for figure's data

## Acknowledgements

Human outbreak strains *Candida auris Charite 607, 611, 588, 610* were obtained from Dr. B. Graf (Charite, Berlin). *Candida albicans 527ΔΔ, Candida albicans 485Δ, Candida albicans MH 203* were provided by Prof. A. Mitchell (University of Georgia, USA).

## Supplementary information 1

Tab.2 Geometrical means, standard deviations and respective confidence intervals. The presentation of values here was done to give statistical information but to prevent overloading the figures of this study with statistical information.

