## Supplementary material for "Comparison of Biofilm quantification in strains of *Candida auris* and *Candida albicans* evaluated by means of Crystal violet, MTT, ATP and NBTZ/BCIP Assays": Sup File 1 Standard deviations and confidence intervals for figure's data: Biofilm MAN-4214684 (1).docx

Jens Jacob

### Robert Koch-Institute, FG16, Seestrasse 10, 13353 Berlin, Germany,

### Corresponding author

**Abstract**

The study presented here shows Biofilm quantification in microtiter plates in strains of *Candida auris* and *Candida albicans* evaluated by means of Crystal violet, MTT, ATP-Luminescence and NBTZ/BCIP assays. The results showed significant differences in biofilm formation between *Candida auris* and *Candida albicans* but also within *Candida auris* outbreak strains in contrast to *Candida auris DSM 21092* reference strain.

**Keywords**: *Candida auris,* biofilm, Crystal violet, MTT, ATP, NBTZ/BCIP

###### Background

Yeasts of the genus *Candida* are members of the *Metschnikowiaceae* and they are phylogenetically closely related to the human non-pathogenic *Saccharomycetacea*. Despite the fact that DNA homology is high between *Saccharomycetacea* and *Metschnikowiaceae*, i.e. only the genera *Candida* within the *Metschnikowiaceae* shows human pathogenic capabilities. Besides the previously known *Candida albicans* there is currently a rise in papers mentioning *Candida auris* as a new emerging human pathogenic yeast [6,16]. *Candida* (and hereby also *C. auris*) are now known worldwide as a cause of human infections, especially in immunocompromised patients. Due to its increasing resistance against antifungal drug [21,23]. However, *Candida* are also able to survive several days under certain environmental conditions especially and foremost in highly regulated biofilms [8]. They also quickly develop resistance against surface disinfectants [7,9,10].

However, in contrast there is both an increasing number of sample submissions to the German consular laboratory for mycoses [NRZ Nationales Referenzzentrum für Invasive Pilzinfektionen (https://www.nrz-myk.de/home.html)](https://www.nrz-myk.de/home.html) [13] and to the German Infectious Diseases Reporting System for infectious disease resulting in a continuous rise of reported *Candida auris* cases (<https://meldung.demis.rki.de/portal/shell/#/welcome>; [https://survstat.rki.de/Default.aspx) [14].](https://survstat.rki.de/Default.aspx)%20%5b14%5d.)

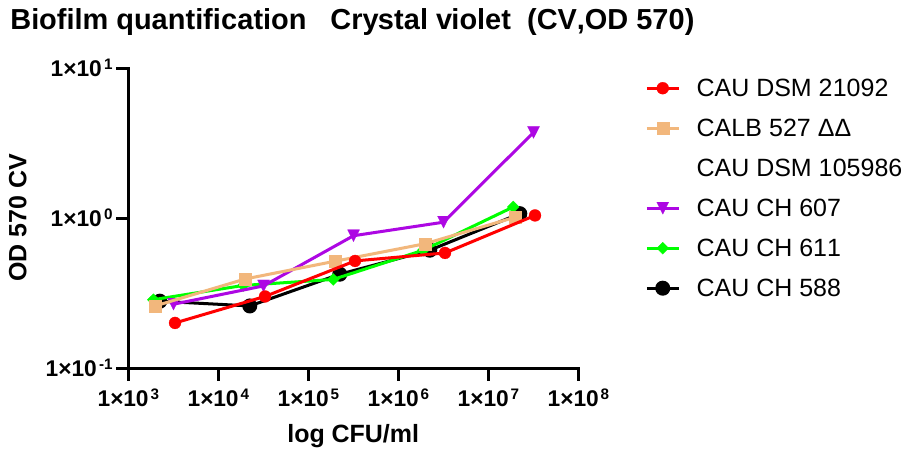

Fig.1. CV OD 570

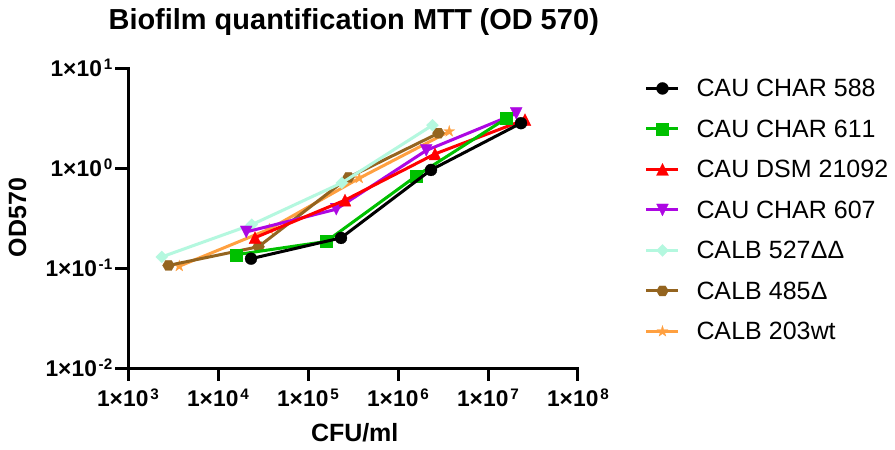

Fig.2. MTT OD 570

**1×10**

**3**

**1×10**

**4**

**1×10**

**5**

**1×10**

**6**

**1×10**

**7**

**1×10**

**8**

**1×10**

**4**

**1×10**

**5**

**1×10**

**6**

**1×10**

**7**

**Biofilm quantification ATP (LUMI 15min)**

**CFU/ml**

**RLU**

CAU CHAR 588 (T 15min)

CAU CHAR 611 (T 15min)

CAU DSM 21092 (T 15 min)

CAU CHAR 607 (T 15 min)

CALB 527ΔΔ (T 15min)

CALB 485Δ (T 15 min)

CALB 203wt (T 15 min)

CAU CH 610 (T 15min)

Fig. 3 ATP Assay

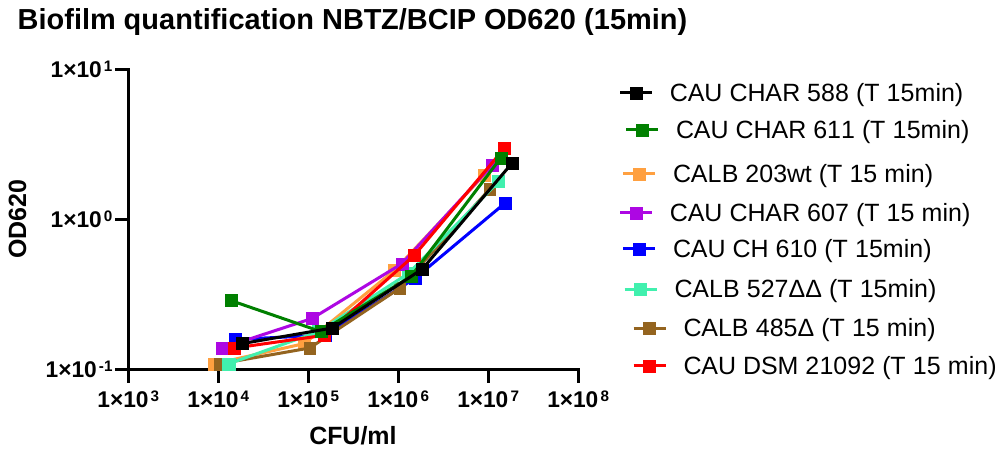

Fig. 4. NBTZ/BCIP Assay (Indigo formation)

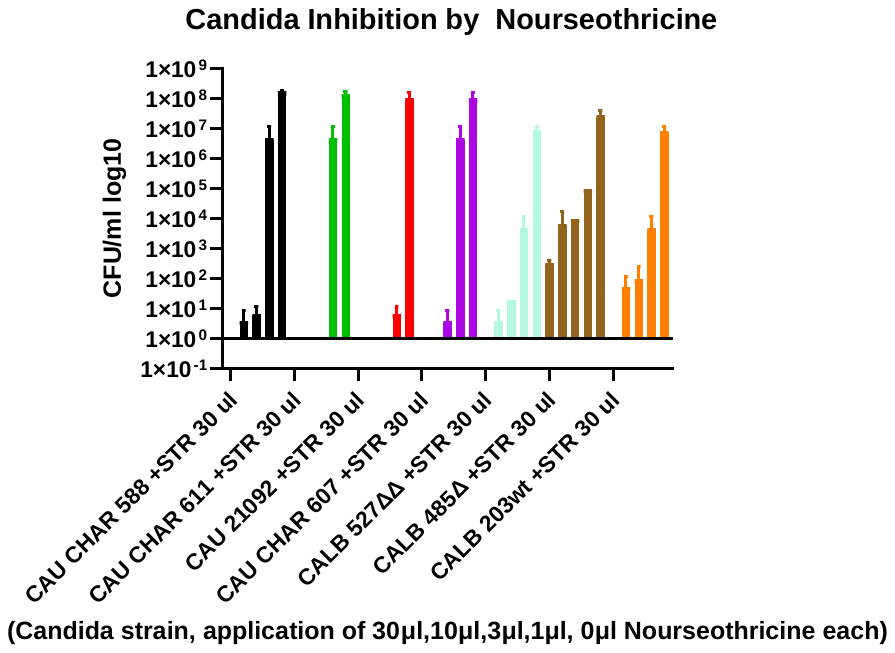

Fig.5 Nourseothricine Inhibition Assay of *Candida* strains.

The (maximal) 5 bars for each organism represents application of 30µl, 10μl, 3µl, 1μl, 0µl Nourseothricine each. All this information would not fit as text in the legend of the y-axis. In Fig. 5 for each strain only STR 30µl is written indicating the first bar of maximum five. If a bar is missing it means no *Candida* growth was measured by plating.

If we have here for example two red bars for strain CAU DSM 21092 only, it means the 30,10 and 3µl Nouseothricine solution inhibited the strains totally (no CFU’s detectable) whereas for 1µl and 0 µl (growth control) growth was visible. According that scheme the graph could be read for all strains.

The description of y axis (30,10,3,1,0 µl Nourseothricine solution in Fig.5, Tab.1) corresponds to300,100,30,10, 0 µg/ml Nourseothricin each.

Tab.1 Nourseothricin inhibition assay

| Candida Strain | Dilution | CFU/ml | ΔOD570- OD 570 (NOUR) Quota | OD 570 (NOUR) | Number of experiments (as technical triplicates each) |
| --- | --- | --- | --- | --- | --- |
| **CAU CH 588** | **1/10** | **10^7^** | **1.2** | **1.6** | **5** |
| **CAU CH 611** | **1/10** | **10^7^** | **1.3** | **1.8** | **5** |
| CALB ΔΔ527 | 1/10 | 10^7^ | 0.6 | 2.2 | 5 |
| **CAU DSM 21092** | **1/10** | **10^7^** | **1.4** | **1.8** | **5** |
| **CAU CH 607** | **1/10** | **10^7^** | **1.6** | **1.8** | **5** |
| CALB Δ485 | 1/10 | 10^7^ | 0.7 | 1.6 | 5 |
| CALB 203wt | 1/10 | 10^7^ | 0.7 | 1.8 | 5 |
| **CAU CH 588** | **1/100** | **10^6^** | **0.5** | **0.4** | **5** |
| **CAU CH 611** | **1/100** | **10^6^** | **0.6** | **0.4** | **5** |
| CALB ΔΔ527 | 1/100 | 10^6^ | 0.1 | 0.6 | 5 |
| **CAU DSM 21092** | **1/100** | **10^6^** | **1.3** | **0.5** | **5** |
| **CAU CH 607** | **1/100** | **10^6^** | **1.0** | **0.4** | **5** |
| CALB Δ485 | 1/100 | 10^6^ | 0.4 | 0.4 | 5 |
| CALB 203wt | 1/100 | 10^6^ | 0.4 | 0.5 | 5 |
| CAU CH 588 | 1/1000 | 10^5^ | 0.2 | 0.2 | 5 |
| CAU CH 611 | 1/1000 | 10^5^ | 0.15 | 0.14 | 5 |
| CALB ΔΔ527 | 1/1000 | 10^5^ | 0.12 | 0.19 | 5 |
| CAU DSM 21092 | 1/1000 | 10^5^ | 0.2 | 0.2 | 5 |
| CAU CH 607 | 1/1000 | 10^5^ | 0.04 | 0.17 | 5 |
| CALB Δ485 | 1/1000 | 10^5^ | 0.02 | 0.15 | 5 |
| CALB 203wt | 1/1000 | 10^5^ | 0.04 | 0.18 | 5 |

Tab. 1 OD 570 MTT Nourseothricin inhibition Assay

OD 570 MTT inhibition Assay. The so obtained OD570 (Nour) values reflects the respective formazan formation in growth inhibited *Candida* cells only. Subtracting these OD570 (Nour) from the non-treated OD 570 values gives an (candida strain specific) quota indicating the relation of metabolism depending formazan formation in contrast to formazan formation in the Nourseothricin growth inhibited candida cells. Quota (strain specific) = OD 570-OD570 (Nour).
