## Supplementary material for "Comparison of Biofilm quantification in strains of *Candida auris* and *Candida albicans* evaluated by means of Crystal violet, MTT, ATP and NBTZ/BCIP Assays": Sup File 1 Standard deviations and confidence intervals for figure's data: Biofilm supplementary.docx

Supplementary File 1: Tab. 2 Descriptive statistic values belonging to Graphs in Fig.’s 1-4

| Fig./subject | Strain/Dilution | Geometric mean | Geometric SD factor | Lower 95% CI of geometric  mean | Upper 95% CI of geometric  mean | Number of biological/technical replicates investigated. |
| --- | --- | --- | --- | --- | --- | --- |
| 1, Crystal violett OD 570 | CAU DSM 21092, 1:10 | 15570 | 2,352 | 10655 | 22751 | 7/21 |
| 1, Crystal violett OD 570 | CAU DSM 21092, 1:100 | 8262 | 2,176 | 5853 | 11662 | 7/21 |
| 1, Crystal violett OD 570 | CAU DSM 21092, 1:1000 | 7161 | 2,203 | 5045 | 10163 | 7/21 |
| 1, Crystal violett OD 570 | CAU DSM 21092, 1:10000 | 4600 | 2,696 | 2892 | 7317 | 7/21 |
| 1, Crystal violett OD 570 | CAU DSM 21092, 1:100000 | 3395 | 2,172 | 2361 | 4880 | 7/21 |
| 1, Crystal violett OD 570 | CALB 527 1:10 | 15019 | 1,722 | 11116 | 20294 | 5/15 |
| 1, Crystal violett OD 570 | CALB 527 1:100 | 7941 | 1,726 | 5869 | 10743 | 5/15 |
| 1, Crystal violett OD 570 | CALB 527 1:1000 | 6592 | 2,144 | 4321 | 10057 | 5/15 |
| 1, Crystal violett OD 570 | CALB 527 1:10000 | 4337 | 2,197 | 2804 | 6706 | 5/15 |
| 1, Crystal violett OD 570 | CALB 527 1:100000 | 3197 | 2,517 | 1917 | 5329 | 5/15 |
| 1, Crystal violett OD 570 | CAU DSM 105986 1:10 | 18324 | 1,430 | 14596 | 23004 | 4/12 |
| 1, Crystal violett OD 570 | CAU DSM 105986 1:100 | 8358 | 1,730 | 5901 | 11838 | 4/12 |
| 1, Crystal violett OD 570 | CAU DSM 105986 1:1000 | 7655 | 1,669 | 5529 | 10598 | 4/12 |
| 1, Crystal violett OD 570 | CAU DSM 105986 1:10000 | 6075 | 1,954 | 3970 | 9298 | 4/12 |
| 1, Crystal violett OD 570 | CAU DSM105986 1:100000 | 3999 | 1,713 | 2841 | 5628 | 4/12 |
| 1, Crystal violett OD 570 | CAU CHAR 607 1:10 | 34726 | 1,823 | 27100 | 44500 | 8/24 |
| 1, Crystal violett OD 570 | CAU CHAR 607 1:100 | 8176 | 2,382 | 5666 | 11796 | 8/24 |
| 1, Crystal violett OD 570 | CAU CHAR 607 1:1000 | 6608 | 2,513 | 4518 | 9666 | 8/24 |
| 1, Crystal violett OD 570 | CAU CHAR 607 1:10000 | 4373 | 2,640 | 2874 | 6655 | 8/24 |
| 1, Crystal violett OD 570 | CAU CHAR 607 1:100000 | 3013 | 2,809 | 1928 | 4709 | 8/24 |
| 1, Crystal violett OD 570 | CAU CHAR 611 1:10 | 15566 | 2,146 | 9014 | 26879 | 4/10 |
| 1, Crystal violett OD 570 | CAU CHAR 611 1:100 | 8490 | 2,487 | 4425 | 16292 | 4/10 |
| 1, Crystal violett OD 570 | CAU CHAR 611 1:1000 | 5940 | 2,659 | 2951 | 11958 | 4/10 |
| 1, Crystal violett OD 570 | CAU CHAR 611 1:10000 | 4424 | 3,929 | 1409 | 13889 | 3/8 |
| 1, Crystal violett OD 570 | CAU CHAR 611 1:100000 | 3603 | 3,045 | 1420 | 9140 | 3/8 |
| 1, Crystal violett OD 570 | CAU CHAR 588 1:10 | 15816 | 2,416 | 8415 | 29725 | 4/10 |
| 1, Crystal violett OD 570 | CAU CHAR 588 1:100 | 8170 | 2,386 | 4386 | 15218 | 4/10 |
| 1, Crystal violett OD 570 | CAU CHAR 588 1:1000 | 5846 | 2,772 | 2819 | 12123 | 4/10 |
| 1, Crystal violett OD 570 | CAU CHAR 588 1:10000 | 3455 | 3,859 | 1117 | 10683 | 3/8 |
| 1, Crystal violett OD 570 | CAU CHAR 588 1:100000 | 3627 | 3,356 | 1318 | 9981 | 3/8 |
| 3, MTT OD 570 | CAU DSM 21092, 1:10 | 3,065 | 1,248 | 2,755 | 3,410 | 6/16 |
| 3, MTT OD 570 | CAU DSM 21092, 1:100 | 1,395 | 1,453 | 1,165 | 1,670 | 6/16 |
| 3, MTT OD 570 | CAU DSM 21092, 1:1000 | 0,4799 | 1,392 | 0,4092 | 0,5628 | 6/16 |
| 3, MTT OD 570 | CAU DSM 21092, 1:10000 | 0,2025 | 1,902 | 0,1455 | 0,2818 | 6/16 |
| 3, MTT OD 570 | CALB ΔΔ527 1:10 | 2,698 | 1,175 | 2,357 | 3,087 | 3/6 |
| 3, MTT OD 570 | CALB ΔΔ527 1:100 | 0,7159 | 1,393 | 0,5427 | 0,9445 | 3/6 |
| 3, MTT OD 570 | CALB ΔΔ527 1:1000 | 0,2725 | 1,584 | 0,1855 | 0,4004 | 3/6 |
| 3, MTT OD 570 | CALB ΔΔ527 1:10000 | 0,1297 | 1,761 | 0,07164 | 0,2349 | 3/6 |
| 3, MTT OD 570 | CALB 203 wt 1:10 | 2,340 | 1,278 | 1,906 | 2,873 | 3/6 |
| 3, MTT OD 570 | CALB 203 wt 1:100 | 0,7954 | 1,310 | 0,6345 | 0,9972 | 3/6 |
| 3, MTT OD 570 | CALB 203 wt 1:1000 | 0,2462 | 1,247 | 0,2047 | 0,2960 | 3/6 |
| 3, MTT OD 570 | CALB 203 wt 1:10000 | 0,1051 | 1,039 | 0,1010 | 0,1094 | 3/6 |
| 3, MTT OD 570 | CAU CHAR 607 1:10 | 3,534 | 1,119 | 3,357 | 3,720 | 6/16 |
| 3, MTT OD 570 | CAU CHAR 607 1:100 | 1,506 | 1,495 | 1,254 | 1,809 | 6/16 |
| 3, MTT OD 570 | CAU CHAR 607 1:1000 | 0,3900 | 1,974 | 0,2861 | 0,5314 | 6/16 |
| 3, MTT OD 570 | CAU CHAR 607 1:10000 | 0,2314 | 1,940 | 0,1646 | 0,3253 | 6/16 |
| 3, MTT OD 570 | CAU CHAR 611 1:10 | 3,151 | 1,235 | 2,816 | 3,526 | 4/11 |
| 3, MTT OD 570 | CAU CHAR 611 1:100 | 0,8408 | 1,403 | 0,7020 | 1,007 | 4/11 |
| 3, MTT OD 570 | CAU CHAR 611 1:1000 | 0,1852 | 1,775 | 0,1364 | 0,2514 | 4/11 |
| 3, MTT OD 570 | CAU CHAR 611 1:10000 | 0,1367 | 1,257 | 0,1198 | 0,1560 | 4/11 |
| 3, MTT OD 570 | CAU CHAR 588 1:10 | 2,827 | 1,185 | 2,582 | 3,094 | 4/11 |
| 3, MTT OD 570 | CAU CHAR 588 1:100 | 0,9650 | 1,465 | 0,7874 | 1,183 | 4/11 |
| 3, MTT OD 570 | CAU CHAR 588 1:1000 | 0,2012 | 1,428 | 0,1664 | 0,2433 | 4/11 |
| 3, MTT OD 570 | CAU CHAR 588 1:10000 | 0,1248 | 1,276 | 0,1084 | 0,1437 | 4/11 |
| 3, MTT OD 570 | CALB Δ485 1:10 | 2,240 | 1,294 | 1,806 | 2,779 | 3/6 |
| 3, MTT OD 570 | CALB Δ485 1:100 | 0,8087 | 1,182 | 0,7032 | 0,9301 | 3/6 |
| 3, MTT OD 570 | CALB Δ485 1:1000 | 0,1643 | 1,311 | 0,1310 | 0,2061 | 3/6 |
| 3, MTT OD 570 | CALB Δ485 1:10000 | 0,1070 | 1,158 | 0,09177 | 0,1248 | 3/6 |
| 4, ATP (LUMI 15 min) | CAU DSM 21092, 1:10 | 1873048 | 3,197 | 857899 | 4089420 | 4/11 |
| 4, ATP (LUMI 15 min) | CAU DSM 21092, 1:100 | 3178258 | 1,810 | 2133660 | 4734272 | 4/11 |
| 4, ATP (LUMI 15 min) | CAU DSM 21092, 1:1000 | 638851 | 1,961 | 406314 | 1004470 | 4/11 |
| 4, ATP (LUMI 15 min) | CAU DSM 21092, 1:10000 | 221577 | 1,429 | 174340 | 281612 | 4/11 |
| 4, ATP (LUMI 15 min) | CALB ΔΔ527 1:10 | 2838974 | 1,859 | 1871793 | 4305909 | 4/11 |
| 4, ATP (LUMI 15 min) | CALB ΔΔ527 1:100 | 2094849 | 1,256 | 1797455 | 2441448 | 4/11 |
| 4, ATP (LUMI 15 min) | CALB ΔΔ527 1:1000 | 316009 | 2,048 | 195230 | 511509 | 4/11 |
| 4, ATP (LUMI 15 min) | CALB ΔΔ527 1:10000 | 245953 | 1,091 | 197977 | 305555 | 4/11 |
| 4, ATP (LUMI 15 min) | CALB 203 wt 1:10 | 3075083 | 1,509 | 2332962 | 4053274 | 4/11 |
| 4, ATP (LUMI 15 min) | CALB 203 wt 1:100 | 1371628 | 1,383 | 1103173 | 1705411 | 4/11 |
| 4, ATP (LUMI 15 min) | CALB 203 wt 1:1000 | 243399 | 1,468 | 188104 | 314947 | 4/11 |
| 4, ATP (LUMI 15 min) | CALB 203 wt 1:10000 | 207030 | 1,100 | 163508 | 262136 | 4/11 |
| 4, ATP (LUMI 15 min) | CAU CHAR 607 1:10 | 3859961 | 1,779 | 2620867 | 5684873 | 4/11 |
| 4, ATP (LUMI 15 min) | CAU CHAR 607 1:100 | 2387150 | 2,179 | 1414404 | 4028897 | 4/11 |
| 4, ATP (LUMI 15 min) | CAU CHAR 607 1:1000 | 437019 | 2,216 | 256060 | 745860 | 4/11 |
| 4, ATP (LUMI 15 min) | CAU CHAR 607 1:10000 | 74499 | 1,967 | 47298 | 117345 | 4/11 |
| 4, ATP (LUMI 15 min) | CAU CHAR 611 1:10 | 4165037 | 2,242 | 2421121 | 7165083 | 4/10 |
| 4, ATP (LUMI 15 min) | CAU CHAR 611 1:100 | 3801614 | 1,364 | 3085683 | 4683654 | 4/10 |
| 4, ATP (LUMI 15 min) | CAU CHAR 611 1:1000 | 468154 | 1,691 | 321539 | 681621 | 4/10 |
| 4, ATP (LUMI 15 min) | CAU CHAR 588 15 1:10 | 5813424 | 1,267 | 4958578 | 6815643 | 4/11 |
| 4, ATP (LUMI 15 min) | CAU CHAR 588 1:100 | 4206976 | 1,206 | 3708539 | 4772404 | 4/11 |
| 4, ATP (LUMI 15 min) | CAU CHAR 588 1:1000 | 426624 | 1,264 | 364421 | 499445 | 4/11 |
| 4, ATP (LUMI 15 min) | CAU CHAR 588 1:10000 | 136541 | 1,795 | 92151 | 202314 | 4/11 |
| 4, ATP (LUMI 15 min) | CALB Δ485 1:10 | 2609916 | 1,275 | 2217042 | 3072410 | 4/11 |
| 4, ATP (LUMI 15 min) | CALB Δ485 1:100 | 1136132 | 1,526 | 855287 | 1509196 | 4/11 |
| 4, ATP (LUMI 15 min) | CALB Δ485 1:1000 | 243518 | 1,859 | 160541 | 369382 | 4/11 |
| 5, NBTZ/BCIP (620 nm, 15 min)-1 | CAU DSM 21092, 1:10 | 2,820 | 1,305 | 2,298 | 3,460 | 3/9 |
| 5, NBTZ/BCIP (620 nm, 15 min)-2 | CAU DSM 21092, 1:100 | 0,5569 | 1,208 | 0,4816 | 0,6439 | 3/9 |
| 5, NBTZ/BCIP (620 nm, 15 min)-3 | CAU DSM 21092, 1:1000 | 0,1644 | 1,222 | 0,1409 | 0,1918 | 3/9 |
| 5 NBTZ/BCIP (620 nm, 15 min)-4 | CAU DSM 21092, 1:10000 | 0,1366 | 1,190 | 0,1195 | 0,1562 | 3/9 |
| 5, NBTZ/BCIP (620 nm, 15 min)-1 | CALB ΔΔ527 1:10 | 1,760 | 1,138 | 1,593 | 1,944 | 3/9 |
| 5, NBTZ/BCIP (620 nm, 15 min)-2 | CALB ΔΔ527 1:100 | 0,4397 | 1,381 | 0,3430 | 0,5635 | 3/9 |
| 5, NBTZ/BCIP (620 nm, 15 min)-3 | CALB ΔΔ527 1:1000 | 0,1805 | 1,428 | 0,1373 | 0,2373 | 3/9 |
| 5, NBTZ/BCIP (620 nm, 15 min)-4 | CALB ΔΔ527 1:10000 | 0,1113 | 1,085 | 0,1045 | 0,1185 | 3/9 |
| 5, NBTZ/BCIP (620 nm, 15 min)-1 | CALB 203 wt 1:10 | 2,043 | 1,060 | 1,954 | 2,137 | 3/9 |
| 5, NBTZ/BCIP (620 nm, 15 min)-2 | CALB 203 wt 1:100 | 0,4619 | 1,186 | 0,4052 | 0,5265 | 3/9 |
| 5, NBTZ/BCIP (620 nm, 15 min)-3 | CALB 203 wt 1:1000 | 0,1504 | 1,253 | 0,1264 | 0,1788 | 3/9 |
| 5, NBTZ/BCIP (620 nm, 15 min)-4 | CALB 203 wt 1:10000 | 0,1081 | 1,156 | 0,09666 | 0,1208 | 3/9 |
| 5, NBTZ/BCIP (620 nm, 15 min)-1 | CAU CHAR 607 1:10 | 2,241 | 1,328 | 1,801 | 2,787 | 3/9 |
| 5, NBTZ/BCIP (620 nm, 15 min)-2 | CAU CHAR 607 1:100 | 0,5060 | 1,169 | 0,4489 | 0,5704 | 3/9 |
| 5, NBTZ/BCIP (620 nm, 15 min)-3 | CAU CHAR 607 1:1000 | 0,2153 | 1,213 | 0,1856 | 0,2498 | 3/9 |
| 5, NBTZ/BCIP (620 nm, 15 min)-4 | CAU CHAR 607 1:10000 | 0,1321 | 1,353 | 0,1047 | 0,1666 | 3/9 |
| 5, NBTZ/BCIP (620 nm, 15 min)-1 | CAU CHAR 611 1:10 | 2,547 | 1,099 | 2,369 | 2,739 | 3/9 |
| 5, NBTZ/BCIP (620 nm, 15 min)-2 | CAU CHAR 611 1:100 | 0,4494 | 1,467 | 0,3348 | 0,6034 | 3/9 |
| 5, NBTZ/BCIP (620 nm, 15 min)-3 | CAU CHAR 611 1:1000 | 0,1856 | 1,231 | 0,1582 | 0,2178 | 3/9 |
| 5, NBTZ/BCIP (620 nm, 15 min)-4 | CAU CHAR 611 1:1000 | 0,1619 | 1,130 | 0,1473 | 0,1778 | 3/9 |
| 5, NBTZ/BCIP (620 nm, 15 min)-1 | CAU CHAR 588 15 min | 2,392 | 1,058 | 2,290 | 2,498 | 3/9 |
| 5, NBTZ/BCIP (620 nm, 15 min)-2 | CAU CHAR 588 15 min | 0,4861 | 1,499 | 0,3561 | 0,6635 | 3/9 |
| 5, NBTZ/BCIP (620 nm, 15 min)-3 | CAU CHAR 588 15 min | 0,1865 | 1,303 | 0,1522 | 0,2286 | 3/9 |
| 5, NBTZ/BCIP (620 nm, 15 min)-4 | CAU CHAR 588 15 min | 0,1478 | 1,238 | 0,1255 | 0,1742 | 3/9 |
| 5, NBTZ/BCIP (620 nm, 15 min)-1 | CALB Δ485 1:10 | 1,558 | 1,148 | 1,401 | 1,732 | 3/9 |
| 5, NBTZ/BCIP (620 nm, 15 min)-2 | CALB Δ485 1:100 | 0,3518 | 1,386 | 0,2736 | 0,4522 | 3/9 |
| 5, NBTZ/BCIP (620 nm, 15 min)-3 | CALB Δ485 1:1000 | 0,1411 | 1,193 | 0,1232 | 0,1615 | 3/9 |
| 5, NBTZ/BCIP (620 nm, 15 min)-4 | CALB Δ485 1:10000 | 0,1102 | 1,109 | 0,1018 | 0,1193 | 3/9 |
| 5, NBTZ/BCIP (620 nm, 15 min)-1 | CAU CHAR 610, 1:10 | 1,303 | 1,144 | 1,175 | 1,446 | 3/9 |
| 5, NBTZ/BCIP (620 nm, 15 min)-2 | CAU CHAR 610, 1:100 | 0,4019 | 1,164 | 0,3576 | 0,4516 | 3/9 |
| 5, NBTZ/BCIP (620 nm, 15 min)-3 | CAU CHAR 610, 1:1000 | 0,1631 | 1,256 | 0,1369 | 0,1943 | 3/9 |
| 5, NBTZ/BCIP (620 nm, 15 min)-4 | CAU CHAR 610, 1:10000 | 0,1517 | 1,271 | 0,1262 | 0,1824 | 3/9 |
